# Non-additive biotic interactions improve predictions of tropical tree growth and impact community size structure

**DOI:** 10.1101/2020.09.16.300616

**Authors:** Hao Ran Lai, Kwek Yan Chong, Alex Thiam Koon Yee, Margaret M. Mayfield, Daniel B. Stouffer

**Affiliations:** Centre for Integrative Ecology, School of Biological Sciences, University of Canterbury, Christchurch 8140, New Zealand; Department of Biological Sciences, National University of Singapore, 16 Science Drive 4, Singapore 117558, Republic of Singapore; Centre for Urban Greenery and Ecology, National Parks Board, Singapore Botanic Gardens, Singapore, Republic of Singapore; School of Biological Sciences, The University of Queensland, St Lucia, QLD 4067, Australia

**Keywords:** Higher-order interaction, Indirect effect, Competition, Facilitation, Diameter growth, Secondary succession, Light limitation

## Abstract

Growth in individual size or biomass is a key demographic component in population models, with wide-ranging applications from quantifying species performance across abiotic or biotic conditions to assessing landscape-level dynamics under global change. In forest ecology, the responses of tree growth to biotic interactions are widely held to be crucial for understanding forest diversity, function, and structure. To date, most studies on plant–plant interactions only examine the additive competitive or facilitative interactions between species pairs; however, there is increasing evidence of non-additive, higher-order interactions (HOIs) impacting species demographic rates. When HOIs are present, the dynamics of a multi-species community cannot be fully understood or accurately predicted solely from pairwise outcomes because of how additional species ‘interfere’ with the direct, pairwise interactions. Such HOIs should be particularly prevalent where species show nonlinear functional responses to resource availability and resource-acquisition traits themselves are density dependent. With this in mind, we used data from a tropical secondary forest—a system that fulfills both of these conditions—to build a ontogenetic diameter-growth model for individuals across ten woody-plant species. We allowed both direct and indirect interactions within communities to influence the species-specific growth parameters in a generalized Lotka–Volterra model. Specifically, indirect interactions entered the model as higher-order quadratic terms, i.e. non-additive effects of conspecific and heterospecific neighbour size on the focal individual’s growth. For the whole community and for four out of ten focal species, the model that included HOIs had more statistical support than the model that included only direct interactions, despite the former containing a far greater number of parameters. HOIs had comparable effect sizes to direct interactions, and tended to further reduce the diameter growth rates of most species beyond what direct interactions had already reduced. In a simulation of successional stand dynamics, the inclusion of HOIs lead to rank swaps in species’ diameter hierarchies, even when community-level size distributions remained qualitatively similar. Our study highlights the implications, and discusses possible mechanisms, of non-additive density dependence in highly diverse and light-competitive tropical forests.

## Introduction

A key pursuit in ecology is to predict the spatio-temporal dynamics of populations (Sutherland et al. 2013). Achieving this goal requires a detailed understanding of the ecological processes that drive species’ demographic performance, such as biotic interactions between species that share resource pools (Tilman 1982). Most studies of resource competition focus on the interactions between species pairs, even when more than two species are involved (Levine et al. 2017). These ‘species pair’ approaches assume that a focal species is simply influenced by the sum of all pairwise interactions between itself and its direct neighbours, although such an additive assumption has long been recognised to likely be a major oversimplification (Abrams 1983; Adler and Morris 1994; Billick and Case 1994; Wootton 1994). Non-additive biotic interactions occur when the direct effect of a neighbour species is modified by other individuals of the same or another species. If this happens, then the strengths of pairwise interactions are no longer constant across communities that vary in composition. When the whole is more than the sum of its parts, even a precise understanding of interactions between species pairs in isolation is insufficient to accurately predict population dynamics in a multispecies assemblage (Billick and Case 1994; Levine et al. 2017; Kleinhesselink, Kraft, and Levine 2019; Letten and Stouffer 2019).

Various biological mechanisms have been proposed for how non-additive biotic interactions may arise. Though their definitions continue to be refined, these mechanisms fall into two general categories: nonlinear density dependence and interaction modification. Both processes have been loosely referred to as higher-order interactions (HOIs), but Kleinhesselink, Kraft, and Levine (2019) recently clarified the distinction referring to them respectively as ‘soft’ and ‘hard’ HOIs. Nonlinear density dependence, or soft HOIs, is the phenomenon when a focal species’ performance does not change linearly with changing neighbour densities, and emerges when species have nonlinear functional responses to resource availability (Letten and Stouffer 2019). Consider, for example, a saturating functional response such as the size growth–light availability relationship in many plants (Rüger et al. 2011; Poorter et al. 2019): at low neighbour density, light resource is plentiful; a small increase in neighbour density thus depletes light at the plateau of the focal individual’s light-response curve, where the competitive impact on size growth is minimal. However, the impact on size growth becomes greater when additional increases in neighbour density deplete light toward the steeper region of the focal’s light-response curve. Soft HOIs also emerge when a neighbour species’ density changes between infrequent sampling events due to *pairwise* interactions, because its effect on the focal species will be apparently different from linear expectations even when the pairwise interaction coefficient remains constant (Kleinhesselink, Kraft, and Levine 2019). Billick and Case (1994) referred to these non-additivities due to continuously changing population densities not captured by discrete-time models as ‘indirect effects’ and draw synonymy to ‘interaction chains’ as defined by Wootton (1993). Importantly, nonlinear density dependence can occur for a focal pair even without a third, intermediary species (Kleinhesselink, Kraft, and Levine 2019; Letten and Stouffer 2019).

Interaction modification, or hard HOIs, on the other hand, arises when a third, intermediary species does not only directly interact with the focal species, but also induces behavioural or plastic changes in the direct-neighbour species, thereby modifying the direct-interaction strength or direction between the focal pair (Wootton 1993; Billick and Case 1994). In multi-trophic systems, for example, the mere presence of a top predator may induce behavioural change in a meso-predator thus modifying the latter’s predation rate on its prey (Adler and Morris 1994). In single-trophic plant–plant interactions, an intermediary species may indirectly influence the focal species by causing plastic change in the direct competitor of the focal species. For instance, the presence of a deep-rooted intermediary species may cause a direct-competitor species to produce shallower roots and hence compete more intensely with a shallow-rooted focal species (Levine et al. 2017). Similarly in a light-limiting forest, an intermediary species just outside of a focal species’ light-interception radius may shade the focal species’ direct neighbours, thereby preventing (or delaying) the direct neighbours from attaining a taller canopy position to shade the focal species. These mechanisms will manifest phenomenologically as non-negligible HOIs, and are expected to be common in systems where resource-acquisition traits such as size are themselves density dependent (Kleinhesselink, Kraft, and Levine 2019). This is because the intermediary species depletes more resources from the focal pair while simultaneously altering the focal pair’s growth. Such a double impact shifts the focal pair’s ability to acquire and deplete each other’s resources thereby modifying their pairwise interaction strength.

Regardless of their mechanistic basis, signals of non-additivity can be statistically detected by fitting quadratic or interaction terms in a phenomenological model (Letten and Stouffer 2019). Recently, Mayfield and Stouffer (2017) presented an analytical framework to quantify and compare direct, pairwise interactions to indirect HOIs through an extension of the phenomenological Lotka–Volterra competition model. The basis of this framework is a regression model that (i) fits the performance of focal species as an additive response to pairwise-interaction effects from direct neighbours and (ii) also includes higher-order quadratic *terms* to allow the strength of these pairwise interactions to be moderated by species outside of or within the focal pair (i.e. introduces density-dependence to the pairwise-interaction coefficients). Note that we follow Kleinhesselink, Kraft, and Levine (2019) to emphasise that higher-order *terms* are distinct from hard higher-order *interactions*. Higher-order *terms* are statistical parameters in our phenomenological model that help capture non-additivities, which do not distinguish the different mechanisms that induce soft and/or hard HOIs from one another. That said, the main purpose of testing if these higher-order quadratic terms are non-zero without sacrificing model parsimony (Pomerantz 1981) is to determine whether or not observed community dynamics can be sufficiently predicted by pairwise interactions alone.

Tropical tree communities naturally meet the conditions under which HOIs are predicted to prevail, yet studies that test for HOIs remain scarce in forest systems. Tropical forests are known for their high primary productivity and biomass accumulation rate during succession (e.g. Poorter et al. 2016), which lead to rapid canopy closure that imposes strong light limitation to the understorey (e.g. Yee et al. 2019). While HOIs can already arise from the nonlinear size-growth response of tree individuals to light extinction due to increasing neighbour densities, the fact that size itself also determines how much light is depleted through shading allows even more room for intermediary species to modify pairwise interactions and give rise to HOIs. Moreover, the relative longevity of perennial trees provides more time for these indirect biotic effects to build up and manifest as detectable HOI signals. In this study, we therefore examine if HOIs are important descriptors of the diameter growth of ten tree species in a tropical secondary forest. While density dependence has received attention in tropical forest studies (e.g. Harms et al. 2000; Comita et al. 2010; Kunstler et al. 2016), the vast majority of them consider only direct interactions. Some studies have incorporated nonlinear density dependence (soft HOIs; e.g. Pacala et al. 1996; Uriarte et al. 2004), but they have not included hard HOIs. With increasing empirical evidence showing pronounced effects of HOIs in herbaceous plant communities (e.g. Weigelt et al. 2007; Mayfield and Stouffer 2017; Xiao et al. 2020) and recently in temperate tree communities (Li et al. 2020), it is becoming important to assess the ubiquity of non-negligible HOIs across more natural systems. If HOIs are widespread and emerge easily under a wide range of conditions (Kleinhesselink, Kraft, and Levine 2019; Letten and Stouffer 2019), their effects likely need to be captured by community models in order to accurately predict the outcome of multi-species interactions both quantitatively (e.g. abundance and size distributions) and qualitatively (e.g. coexistence vs. competitive exclusion; Levine et al. 2017). We expect HOIs to emerge in our tropical forest system and, if so, to add to the growing empirical evidence for HOIs in annual and perennial plant communities.

## Methods

### Data collection

The community data originated from Yee et al. (2019) who surveyed a secondary lowland tropical forest in the Central Catchment Nature Reserve, Republic of Singapore (known locally as the ‘Mandai forest’; 1°24.8′N, 103°47.5′E). Regenerating for at least 80 years, the Mandai forest is a mixture of young and old secondary forest patches characterised by both early- and late-successional native plant species. The climate is tropical with annual precipitation of 1,300–2,700 mm yr^−1^ and mean daily temperatures of 26–29°C across the study period. Yee et al. (2019) originally designed the study to track the recovery of woody plant communities from a windstorm disturbance on 11 February 2011. Within 3 months following the windstorm, forty 10 × 10 m^2^ plots were established randomly in blowdown areas with a minimum 40 m distance between plots. Five annual censuses were conducted between April and August in 2011–2015, during which we counted, identified and measured the diameter-at-breast-height (DBH, cm) of all woody stems ≥ 1 cm DBH in each plot.

For this study, we selected the 10 focal tree species that were the most common species by abundance and provided sufficient data for the analyses that follow. These species naturally span a range of slow-to-fast diameter growth and included *Archidendron clypearia* (Jack) I. C. Nielsen, *Calophyllum wallichianum* Planch. & Triana var. *incrassatum*, *Elaeocarpus mastersii* King, *Garcinia parvifolia* Miq., *Gironniera nervosa* Planch., *Macaranga bancana* (Miq.) Mull. Arg., *Palaquium obovatum* (Griff.) Engl., *Prunus polystachya* (Hook. f.) Kalkm., *Syzygium borneense* (Miq.) Miq., and *Timonius wallichianus* (Korth.) Valeton. Total abundances of each focal species range from 116 to 956 (median = 287) giving a total of 3,268 observations (see Table S1 in Supplementary Information).

### Analyses

We calculated the absolute growth rate, 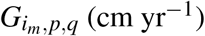, of focal individual *m* of species *i* observed in plot *p* and year *q* as the increment in diameter, *D* (cm), between the census intervals, *t* (yr): 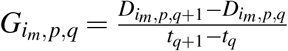, and then modelled growth as a function of diameter using the size-decline growth equation (Zeide 1993, see also Chong et al. 2017),

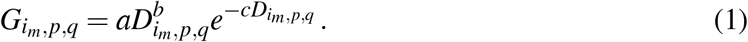

In Equation 1, parameter *a* mainly determines the initial growth rate at small diameters; 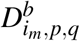 is the ‘size expansion’ component of diameter growth where *b* describes the power relationship between absolute growth rate and size; and *c* describes the exponential decline in absolute growth rate with size due to various physiological limitations. Importantly, we included the ‘size decline’ component, 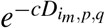, instead of fitting a simple power law function 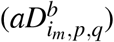 to improve parameter estimation by preventing the biotic interaction coefficients (see *α*’s and *β*’s below) from compensating for reductions in size growth due to *internal* causes, such as metabolic costs of being large. As all of *a*, *b*, and *c* are expected to be positive and non-zero, Equation 1 qualitatively predicts absolute growth rate to increase with diameter at smaller sizes followed by a decline as size increases. This results in the hump-shaped growth–diameter relationship observed in many tree species and forest systems (e.g. Kunstler et al. 2016) including ours.

Although Equation 1 is intended for non-zero, positive growth values, we included negative and zero growths because they constituted 24% of observations. Removing them could overestimate growth or lose information about biotic interactions. Therefore, we assumed absolute growth rate 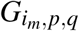 followed a Gaussian distribution with mean 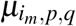 and variance 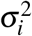. While mean diameter growth 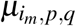 remains constrained to the positives, the variance 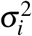 allows non-positive growths to realise due to stem shrinkage or measurement errors. To fulfil the Gaussian assumption, we followed Condit et al. (2017) and performed a modulus transformation on 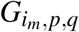 to reign in the right-skewed positive and left-skewed negative values:

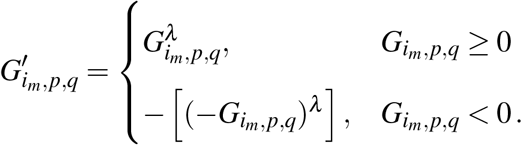

We used the power *λ* = 0.55 (also within the range of Condit et al. 2017) because it gave transformed growth, 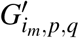 with the lowest skewness. The statistical model is thus:

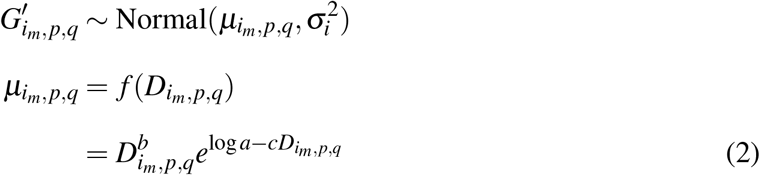

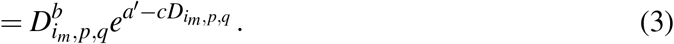

Note that we reparameterised Equation 1 to 2 and then 3 by defining the logarithmic initial growth rate at small diameters, *a*′ = log *a*, so that the three growth parameters *a*′, *b*, and *c* had more similar scales which assisted model convergence.

The diameter–growth model was originally intended to be fit to data from a single species. To accommodate our community data pooled across multiple species, we expanded Equation 3 under the multilevel modelling framework such that each of the growth parameters (*a*′, *b*, and *c*) are partitioned into population-level estimates (‘fixed effects’) and multiple species-specific estimates (‘random effects’). In our multilevel model, *a*′, *b*, and *c* are then estimated to respectively be 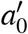, *b*_0_, and *c*_0_ on average while varying by 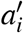, *b_i_*, and *c_i_* for species *i*. To further account for spatiotemporal variations in diameter growth, we included both plot-specific and year-specific effects— 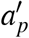 and 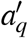 —on the average logarithmic growth rate *a*′, such that:

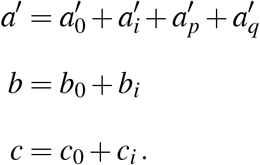

Incorporating plot-specific ‘random’ effects also helps to account for differences in edge effects in our spatially-implicit model. Equation 3 hence becomes:

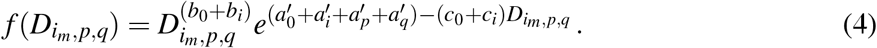

To incorporate biotic interactions into the diameter–growth equation, we first modified Equation 4 to include the cumulative effect of direct interactions with neighbouring species, *g*(*A_j,p,q_*) on 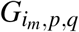 in a generalised Lotka–Volterra fashion:

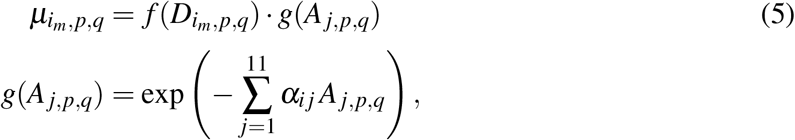

where *A_j,p,q_* is the total basal area (cm^2^) of neighbour species *j* in plot *p* and year *q*, and *α_i j_* are pairwise interaction coefficients that quantify the per-basal-area direct effects of species *j* on growth of the focal species *i*. Note that we included all neighbour individuals in the calculation of *A_j,p,q_*. We generalised the Lotka–Volterra form in Equation 5 such that *α_i j_* can be positive or negative to encompass both competitive and facilitative interactions. The cumulative proportional effect of direct interactions on focal species *i* is then the sum of *α_i j_A_j,p,q_* across all neighbour species. When *i* = *j*, *α_i j_* = *α_ii_* is the measure of intraspecific direct interaction. Note that there are eleven instead of ten neighbour species (*j* = 1,2,…,11) because we included the total basal areas of all remaining non-focal species in each community as an eleventh neighbour group (Martyn et al. 2021). Including these other species helps to minimise the chance of falsely concluding the presence of higher-order interactions when inaccurate growth prediction could stem from unaccounted direct interactions (Billick and Case 1994).

We next incorporated cumulative effects of indirect, higher-order interactions (HOIs) amongst species into Equation 5 following Mayfield and Stouffer (2017):

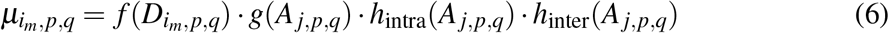

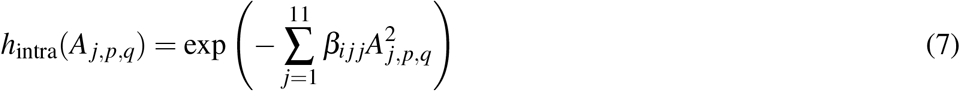

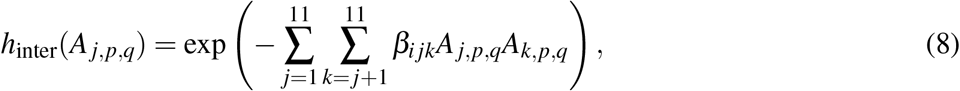

where *β_ijj_* captures the higher-order (i.e. quadratic) effect of neighbour *j* on the direct interaction between species *j* and focal species *i*, and is hereafter coined ‘intraspecific HOI’ (after Mayfield and Stouffer 2017) as this higher-order term takes place between conspecifics of species *j*. On the other hand, *β_ijk_* captures the higher-order effect of a heterospecific neighbour *k*’s total basal area, *A_k,p,q_*, on the direct interaction between neighbour species *j* and focal species *i*, and is hereafter coined ‘interspecific HOI’. In Kleinhesselink, Kraft, and Levine (2019), the intraspecific HOI terms, *β_ijj_*, were referred to as soft HOIs since they still only involve the directly interacting species pair. This helps distinguish them from the interspecific hard HOI interactions, *β_ijk_*, that involve a third species that could modify how the first two species interact in a multi-species community. In this study, we include both soft and hard higher-order *interactions* as non-additive *terms* in the HOI-inclusive model as they all capture non-additivities in any neighbour species’ biotic influence over the focal species. Alternatively, Equation 7 and 8 can be written as:

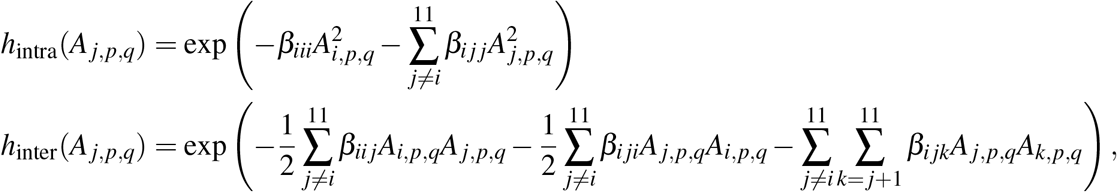

so that the intra- and interspecific direct interaction coefficients can each be compared to their HOI counterparts: namely, (i) *α_ii_* with *β_iii_* and 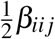 and (ii) *α_ij_* with 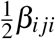, *β_ijj_*, and *β_ijk_*.

### Model fitting

Prior to model fitting and to assist model convergence, we standardised 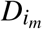 to unit standard deviation and also normalised and standardised *A_j_* to a mean of zero and unit standard deviation. We fit three models in total (Equations 4, 5, and 6) and estimated the parameters through Bayesian inference by fitting non-linear hierarchical models in Stan (Stan Development Team 2018) using the `brm` function in the `brms` package (Bürkner 2017) in `R`. For the population-level ‘fixed’ parameters, we used a weakly informative Normal(0, 10) prior for 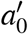 and Halfnormal(0, 10) priors for both *b*_0_ and *c*_0_. For the standard deviations of group-level ‘random’ parameters (i.e. all parameters with subscript *i*, *p*, or *q*, including *α* and *β*), we used a weakly informative Student-*t* prior with 3 degrees of freedom, zero mean and one standard deviation. The parameter posterior distributions were obtained after four chains of 3,000 Hamiltonian Monte Carlo (HMC) warmup iterations followed by 1,000 HMC sampling iterations. We considered models as converged when the 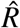 value of all parameters across chains were < 1.05 (Vehtari et al. 2019).

### Model comparison

To assess if the inclusion of direct and/or higher-order interaction terms are necessary for a parsimonious explanation of diameter growth, we compared the null, direct-interaction-only, and HOI-inclusive models (i.e. Equations 4, 5, and 6, respectively) using three goodness-of-fit measures: Bayes *R*^2^, Widely Applicable Information Criteria (*WAIC*), and Leave-One-Out cross-validation Information Criteria (*LOOIC*). These goodness-of-fit measures were chosen to complement one another: Bayes *R*^2^ quantifies the expected fit or variance explained by a model; both *WAIC* and *LOOIC* also measure the expected fit of a model, but they penalise a model with a greater number of effective parameters (‘overfitted’) and hence predicts poorly out-of-sample. *LOOIC* also provides additional checks against *WAIC* because the former is more robust against weak priors and influential data (Vehtari, Gelman, and Gabry 2017). Bayes *R*^2^, *WAIC*, and *LOOIC* were computed for the whole dataset, as well as separately for each focal species. Because both *WAIC* and *LOOIC* are sums across observations and hence increase with sample size, they need to be standardised to a fixed number of observations for a fair comparison among focal species that varied in sample size. Therefore, for each species we additionally bootstrapped its observations with replacement to *n* = 116, which is the lowest number of observations among the 10 focal species (Table S1). We performed this resampling 1,000 times and obtained the median *WAIC* and *LOOIC* with distribution percentiles at *n* = 116.

### Simulation

The above analysis examines variation in instantaneous annual diameter growth rate. We additionally explored how the exclusion or inclusion of HOI in modelling would influence predicted forest-stand structure and community dynamics over a longer timeframe. To do so, we used both the direct-interaction-only and HOI-inclusive models to numerically simulate the temporal change in diameter for each focal species growing under three recruitment scenarios (see below). All simulations were assumed to take place under an average spatiotemporal condition, so both plot and year effects (*a_p_* and *a_q_*) were set to zero in each time step.

To have a realistic initial neighbourhood composition, we used a joint species distribution model published from the same study site (Lai et al. 2020) to predict the recruitment of each focal species in a given plot under 100% canopy openness and other environmental variables (i.e. leaf litter depth, soil nitrogen, phosphorous, potassium, and forest type) at their averages in the first census since wind disturbance. We specifically compared ‘low’, ‘median’, and ‘high’ initial recruitment scenarios, which correspond to the 5th, 50th, and 95th percentiles of predicted recruitments. Because the Lai et al. (2020) model also predicts that recruitment of other non-focal species would constitute roughly half (50.9%) of the total recruitment across species in a plot, we replaced these non-focal recruits with our focal recruits by doubling the predicted focal recruitments to obtain the focal species’ initial abundances; this resulted in 16, 26, and 46 initial stems for the ‘low’, ‘median’, and ‘high’ initial recruitment scenarios, respectively (Table S2). We assumed all recruits begin at 1-cm DBH and then used parameters inferred from both the ‘direct-interaction-only’ model (Equation 5) and the ‘HOI-inclusive’ model (Equation 6) to simulate individual diameter growth of focal species under the three recruitment scenarios at daily timesteps over two years—the time taken for canopy closure in our study site (Yee et al. 2019).

## Results

Compared to both the null and direct-interaction-only models, the HOI-inclusive model had a greater predictive accuracy across the pooled or species-specific observations, as judged by the Bayes *R*^2^ (Fig. 1a). A greater *R*^2^ is not surprising given that the HOI-inclusive model has approximately 90 more effective parameters. Nevertheless, the HOI-inclusive model was still judged by both *WAIC* and *LOOIC* as a far better model for the pooled data (Figs 1b and 1c). When *WAIC* and *LOOIC* were resampled and calculated for each species separately, the HOI-inclusive model performed better than both the null and direct-interaction-only models (Δ*WAIC < −*2 and Δ*LOOIC < −*2) for four out of the ten focal species as well as for all focal species pooled, but performed as well or worse for the other six focal species.

**Figure 1:**
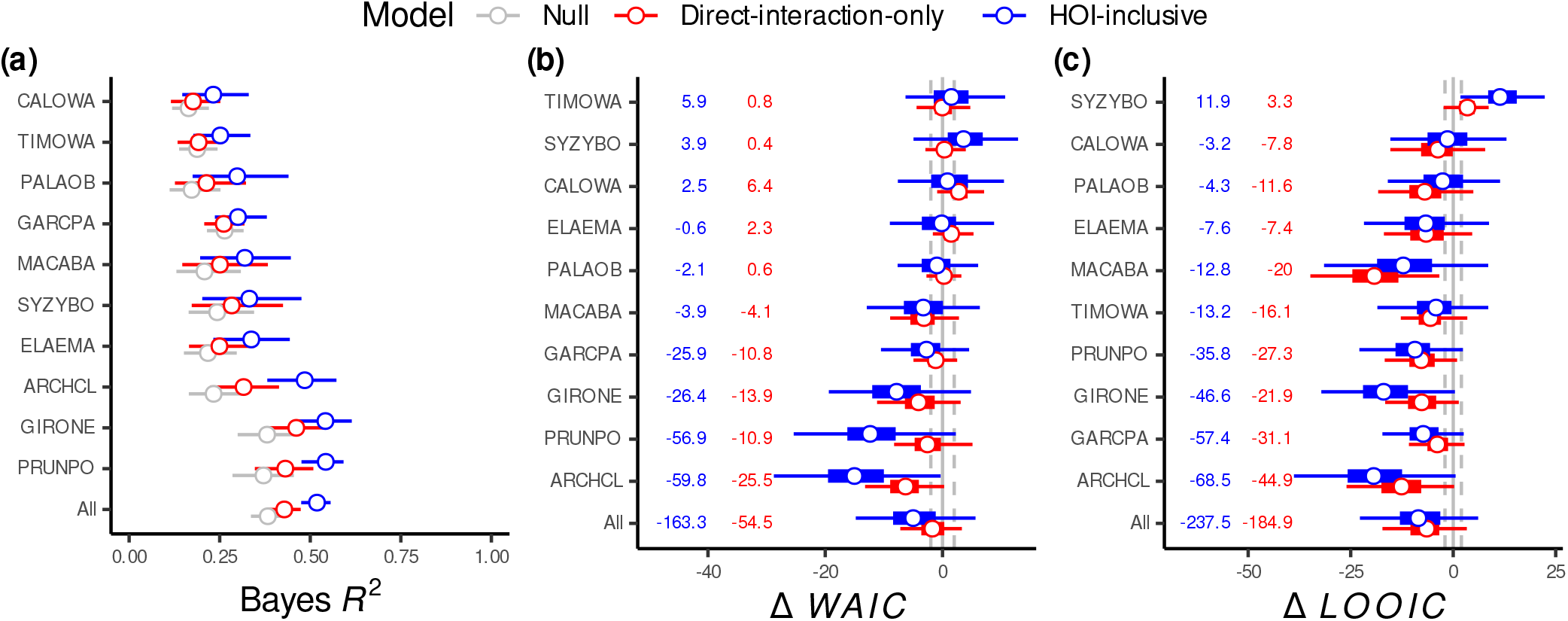
Comparing the goodness-of-fit among the null model (grey), direct-interaction-only model (red), and HOI-inclusive model (blue) in terms of **(a)** Bayes *R*^2^, **(b)** difference in *WAIC*, and **(c)** difference in *LOOIC* for each focal species or all focal species pooled. In **(a)**, circles and horizontal bars denote the median and 95% credible intervals of Bayes *R*^2^, respectively. In **(b)** and **(c)** respectively, circles, thick and thin horizontal bars denote the median, 50% and 95%-tile intervals of the resampled Δ*WAIC* and Δ*LOOIC* to *n* = 116 of both direct-interaction-only (red) and HOI-inclusive models (blue) compared to the null model, while vertical dashed lines denote ±2 Δ*WAIC* or Δ*LOOIC* from zero. The numbers adjacent to each species are the actual Δ*WAIC* or Δ*LOOIC* summed across all observations without resampling. See Table S1 for key to species abbreviations.

The standardised HOI coefficients (*β*’s) among focal species have magnitudes that are comparable to those of direct interactions (*α*’s; Fig. 2). The medians of most direct-interaction coefficients between conspecifics (*α_ii_*’s; 70%) and their corresponding HOI coefficients (*β_ii_.*’s; 64%) have positive signs, i.e. competitive (Fig. 2a). More than half of the medians of the interspecific direct interactions (*α_ij_*’s; 58%) and their corresponding HOI coefficients (*β_ij_.*’s; 70%) are competitive (Fig. 2b). Overall, 35% of these interaction coefficients have negative medians, i.e. facilitative. The interaction coefficients involving non-focal neighbour species also have similar magnitudes and tendencies to be positive compared to that of the focal species (Fig. S1). In most observed cases, however, these interaction coefficients manifested as small effects with 90% of focal individuals experiencing 0.94–1.01 proportional change in annual diameter growth rates due to an *individual* neighbour tree (i.e. growth rate was reduced *multiplicative* to 94% or slightly increased to 101% of its maximum value; Fig. S2a).

**Figure 2:**
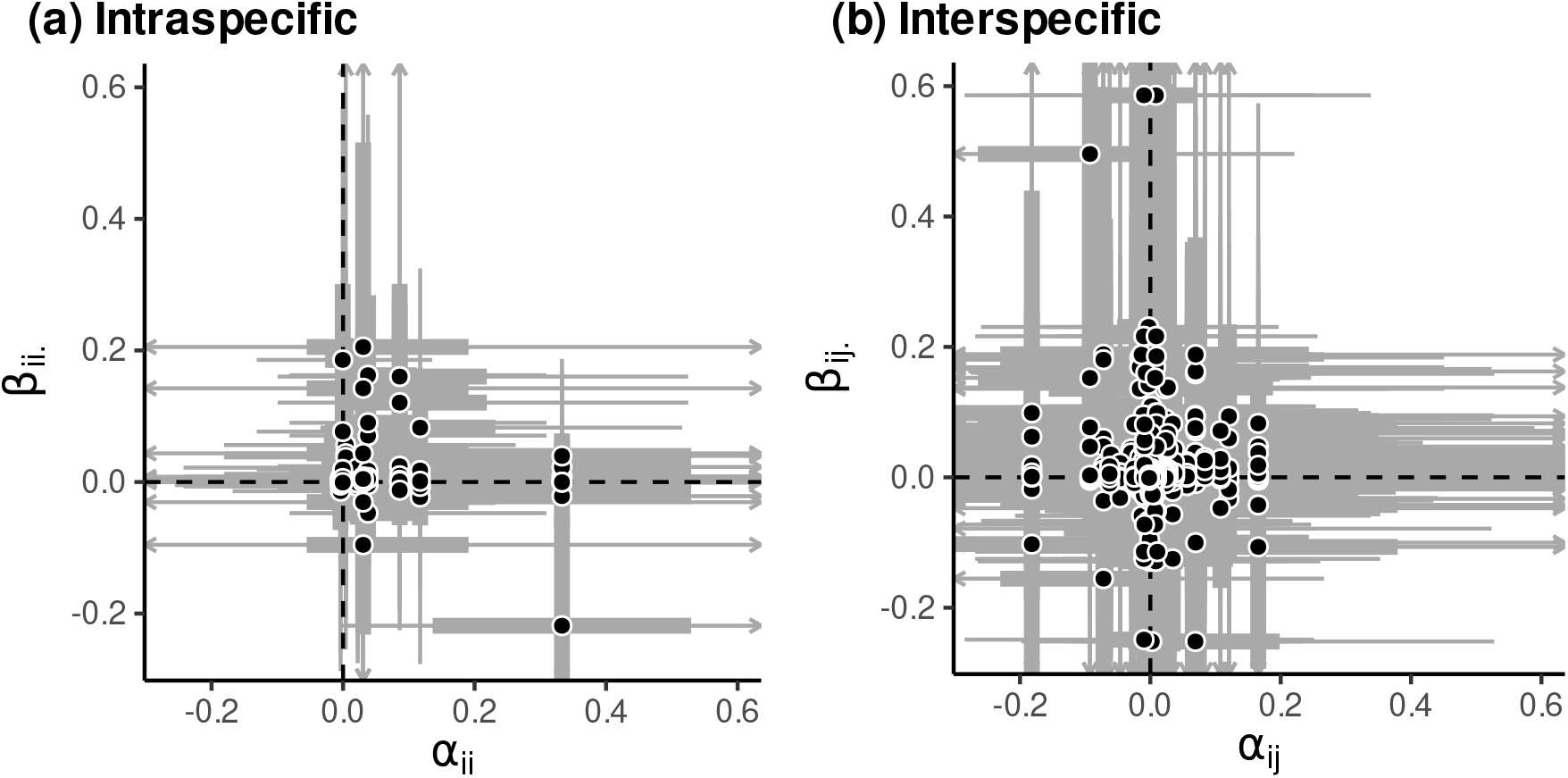
The relationship between direct-interaction coefficients (*α*’s) and HOI coefficients (*β*’s). Both axes are standardised coefficients that have comparable magnitudes. In **(a)**, intraspecific direct-interaction coefficients (*α_ii_*) are plotted with their corresponding HOI coefficients (*β_iii_* or 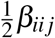, together denoted *β_ii_.*). Similarly in **(b)**, interspecific direct-interaction coefficients (*α_ij_*) are plotted with their corresponding HOI coefficients (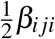, *β_ijj_* or *β_ijk_*, together denoted *β_ij_.*). Points are median estimates with 50% and 95%-tile intervals across the posteriors (thick and thin bars; arrows denote 95%-tile intervals that extend beyond the plot limits).

When the effects of individual neighbour trees are compounded in a multispecies assemblage, these direct interactions and HOIs cumulatively reduced focal species’ annual diameter growth rates by varying magnitudes (Figs 3 and 4). Under the average neighbour basal areas, direct interactions reduced the diameter growth rates of eight out of ten focal species, whereas HOIs always further reduced diameter growth rates (Fig. 3). This resulted in 90% of focal individuals having their observed diameter growth fall between one-third (34%) and slightly above (103%) of their average potential growth rates due to all biotic interactions in combination (Fig. S2b). In absolute terms, this translates to a median reduction in peak growth of −0.09 cm yr^−1^ for the slowest-growing species, *Calophyllum wallichianum* var. *incrassatum* (labelled CALOWA), up to −6.53 cm yr^−1^ for the fastest-growing species, *Prunus polystachya* (PRUNPO; Fig. 3). The median cumulative effects of intraspecific direct interaction, 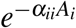, which were multiplicative ranged between 0.86–1.00 across focal species (X axis in Fig. 4a). The median cumulative effects of the HOIs, 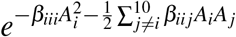, which modify intraspecific direct interactions had a similar range of 0.85–1.00 (Y axis in Fig. 4a). On the other hand, the median cumulative effects of interspecific direct interactions, 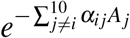 had a narrower range (0.97–1.04) than their corresponding interaction modifiers, 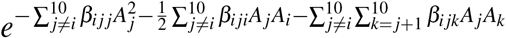 that lay between 0.85–0.98 (Fig. 4b). Combined, there is a weak negative association between the median cumulative effect of all HOIs, 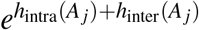 (ranged 0.71–0.96) and that of all direct interactions, 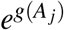 (ranged 0.83–1.04; Fig. 4c).

**Figure 3:**
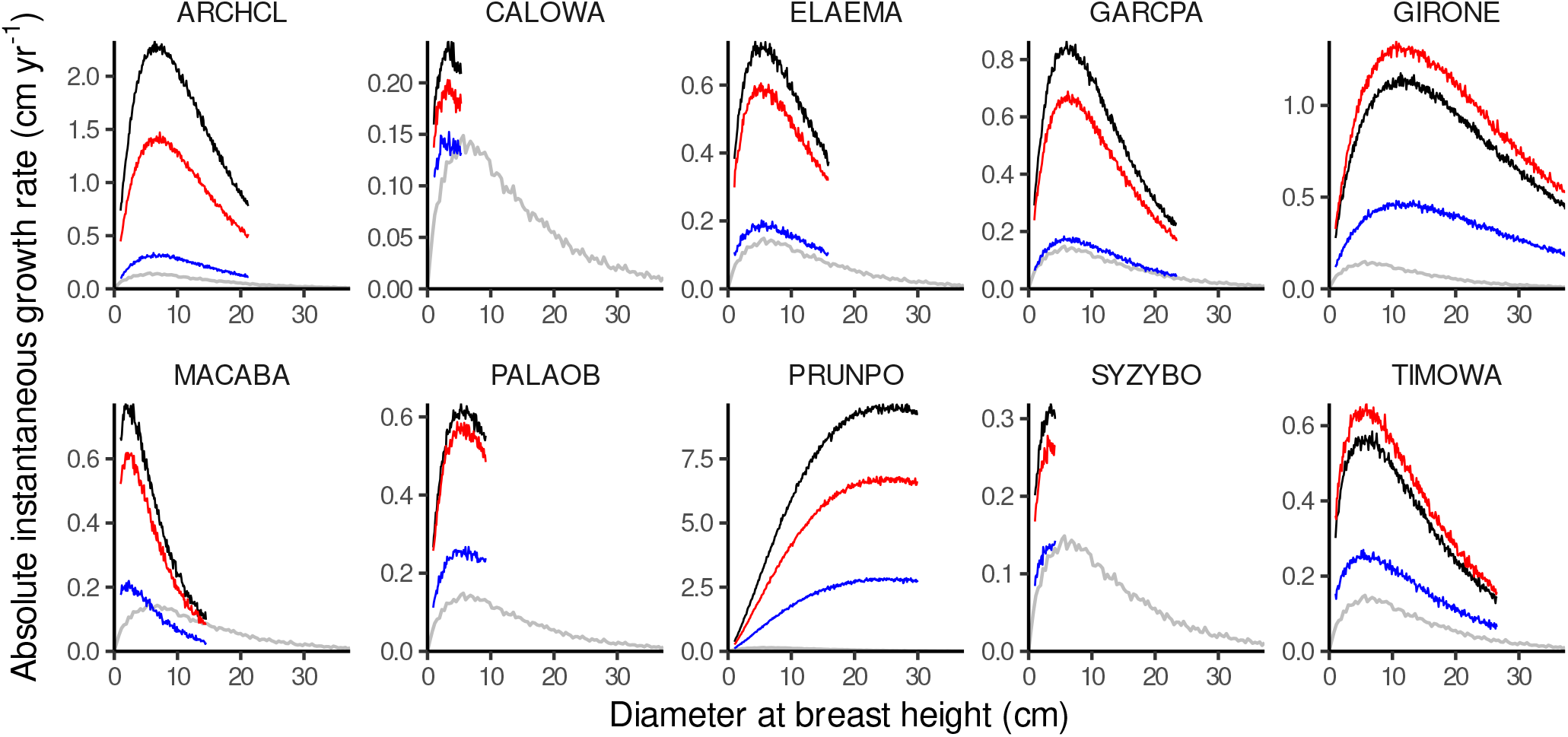
Median of absolute instantaneous growth rate of focal species, 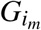, with increasing diameter-at-breast-height (DBH, *D*; cm) under three combinations of biotic interaction terms: no biotic interactions (black curves), direct interactions only (red), and all biotic interactions including HOIs (blue). For the two latter scenarios, predictions were made with all neighbours set at their average total basal areas at any space and time. For a baseline for comparison, grey curves show the absolute growth rate of an ‘average’ species (prediction without species-specific ‘random’ effects) when all types of biotic interaction are taken into account. Diameter growth values were modulus-transformed prior to analyses but are here back-transformed to their original scale. Note the different scales on the Y-axes. The diameter ranges in each panel have been truncated to cover each species’ observed range.

**Figure 4:**
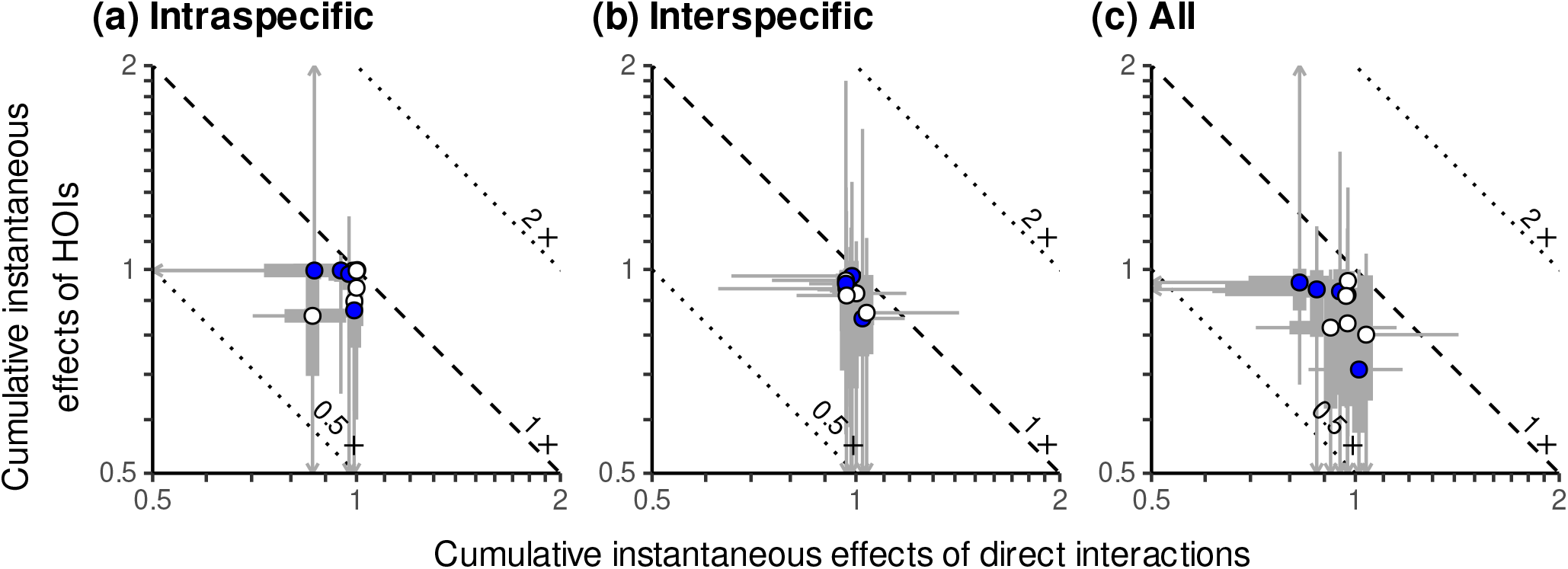
Comparing the observed cumulative effects of direct-interactions and HOIs on the absolute instantaneous growth rate of a focal species. In **(a)**, the cumulative effects of intraspecific direct interactions, 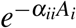, are compared against their corresponding cumulative HOI effects, 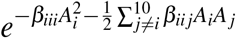. In **(b)**, the cumulative effects of interspecific direct interactions, 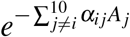, are compared against their corresponding cumulative HOI effects, 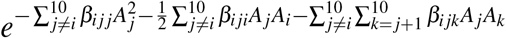. In **(c)**, the cumulative effects of all direct interactions, 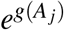 are compared against the cumulative effects of all HOIs, 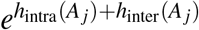. Both axes represent the proportional change in the average absolute diameter growth rate of a focal individual: a focal individual experiences decreases in growth rate when 0 < cumulative effects < 1 (‘competitive effect’) but increases in growth rate when cumulative effects > 1 (‘facilitative effect’). Each circle represents a focal species’ median with 50% and 95%-tile intervals across individuals (thick and thin bars; arrows denote 95%-tile intervals that extend beyond the plot limits). Blue filled circles are the four focal species with the HOI-inclusive model as the best supported as judged by *WAIC* (see Fig. 1b). The dashed diagonal line denotes an isocline where there is no *total* proportional change in the absolute growth rate due to the total cumulative effect of both axes cancelling each other out, i.e. values of both axes multiply to 1. The lower and upper dotted diagonal lines denote a total cumulative effect that halves and doubles the absolute growth rate, respectively. The diameter ranges in each panel have been truncated to cover each species’ observed range. Note the log-scale on both axes.

In addition to the aforementioned *instantaneous* effect of HOIs on focal species’ diameter growth, we examined the *short-term* effect of HOIs by simulating the ten focal species growing together over two years under low, median, and high recruitment scenarios, and then compared the community size structures resulting from the direct-interaction-only and the HOI-inclusive models (Fig. 5). Between the two models, there was no qualitative difference in community size structure at the end of the second year across the three recruitment scenarios (Figs 5a and b). However, there were clear rank swaps in the size hierarchy: the species identity of individuals that grew most and least changed when HOIs were taken into account (Fig. 5c). Overall, rank swaps in the final diameter between models occurred with greater magnitudes as the initial recruitment decreased, even for the focal species whose HOI-inclusive model was not the best model as judged by either *WAIC* or *LOOIC*. Focal species varied in the direction, magnitude, and consistency of rank swap across recruitment scenarios. When HOIs were included, the top-three fastest growing species (with the highest 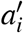 in Equation 4)—*P. polystachya* (PRUNPO), *Macaranga bancana* (MACABA), and *Gironniera nervosa* (GIRONE)— ranked lower under the low recruitment scenario, but their rank reductions were mitigated under median and high recruitment densities. In contrast, the slower-growing species tend to have increased or similar size rank in the presence of HOIs, with the slow-growth *C. wallichianum* var. *incrassatum* and intermediate-growth *Palaquium obovatum* (PALAOB) having the most consistent positive rank swap when HOIs were included.

**Figure 5:**
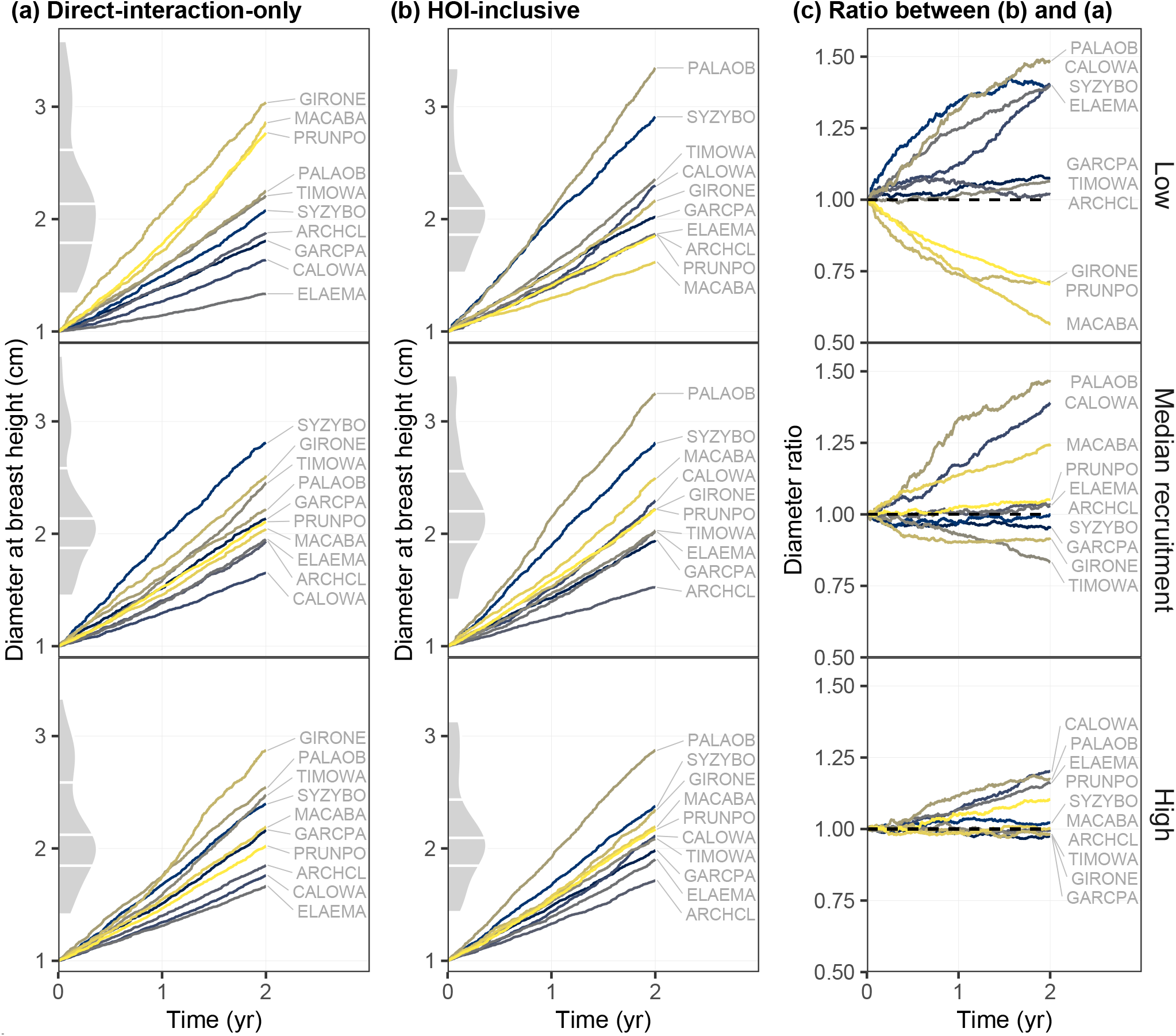
Two-year diameter-growth simulations of the ten focal species growing under low, median, and high recruitment scenarios: **(a)** predictions from the direct-interaction-only, **(b)** predictions from the HOI-inclusive model, and **(c)** the ratio of simulated diameter between HOI-inclusive and direct-interaction-only models. Each line represent an focal species colour-coded by yellow to blue representing increasing initial growth rates at small diameters, 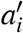 (Equation 4). Though some species may have multiple individual recruits (Table S2), each line is the median per species for visual clarity (see Fig. S3 for individual-tree lines). Density plots with quartile lines along the Y-axes show the size distribution of individuals trees at year two.

## Discussion

Building on early literature that questioned the consequences of ignoring non-additive biotic interactions (e.g. Neill 1974; Abrams 1980; Wootton 1994), recent empirical work has provided evidence for non-negligible higher-order interactions (HOIs) in various natural systems (e.g. Weigelt et al. 2007; Mayfield and Stouffer 2017; Li et al. 2020; Xiao et al. 2020) and prompted theoretical research into the conditions under which HOIs should be expected to emerge (Kleinhesselink, Kraft, and Levine 2019; Letten and Stouffer 2019). Tropical forests meet two of the proposed conditions for emergent HOIs: (i) resource acquisition traits, such as size, that are themselves density-dependent, and (ii) growth in size that responds non-linearly to resource availability. Using a tropical secondary forest dataset from Singapore, we found that the inclusion of HOIs improved the prediction of tree diameter growth for the whole community and for at least four out of ten focal tree species at the species level in this dataset. The inferred HOIs have comparable standardised effect sizes to direct interactions, and tend to further reduce diameter growth rates beyond what direct interactions had already reduced. Even for the other focal species that were less sensitive to the instantaneous effect of HOIs, HOIs could still influence their diameter growth rate by suppressing their competitors’ or facilitators’ size over a longer period of time (as discussed below).

Although our phenomenological model does not mechanistically pinpoints the exact indirect processes leading to HOIs, it adds to the accumulating empirical support for HOIs by demonstrating the presence of non-additive density dependencies in perennial plant systems. That said, mechanisms that gave rise to the detected non-additivities in our study site were likely a mix of nonlinear density dependencies and interaction modifications. As mentioned earlier, these mechanisms include the nonlinear functional response of size growth to light availability (Kleinhesselink, Kraft, and Levine 2019). As neighbour basal areas build up during succession, the strength of biotic interactions change with accelerating or decelerating rates depending on where the forest stand is along the light availability gradient. Such indirect effects due to changing neighbour densities (Billick and Case 1994) are likely given the rapid diameter growth of some focal species under some conditions (greater than the 1-cm yr^−1^ cutoff that defines fast growth in Rüger et al. 2018). Between our annual censuses, these fast-growing neighbours could reach large sizes and reduce (or encourage) the growth of neighbours, thereby reducing (or increasing) the latter’s effect on a focal individual through an interaction chain (Levine et al. 2017). Alternatively, non-additivities in our study could arise from interaction modification when an intermediary species induces plastic changes in the direct neighbour’s morpho-physiological traits, such as stem inclination or side-branching due to phototropism or a narrower crown due to investment in a more slender stem to overcome shading (Sterck, Bongers, and Newbery 2001; Iida et al. 2012), thus casting more or less shade on the focal individual. If this architectural change is long-term, then the modified pairwise interaction will persist even after the intermediary species’ basal area stopped changing. The list of mechanisms here may include an explanation as to why our HOIs exacerbated, rather than mitigated (as in Li et al. 2020), the reduction in diameter growth rates attributable to direct interactions.

More controlled setups are required to conclusively identify mechanisms, but it is clear from our results that HOIs are often crucial at least for the purpose of accurate prediction. At the species level, the HOI-inclusive model more accurately predicted the diameter growth for four out of ten focal species. Though this may seem to indicate that many other species are insensitive to HOIs, we argue that HOIs should not necessarily be considered unimportant for these species in a community-wide context. For observational data, the results for any given focal species can depend as much on what that species is as with which neighbours it happens to be next to. As a result, there are contexts in which a species that is more sensitive to direct than higher-order interactions can still be affected by HOIs; for example, when it is responding to an HOI-sensitive direct neighbour. This could be especially prevalent when this type of neighbour species is locally common or attains a very large size. In our short-term simulation, for example, *M. bancana* (MACABA) was a focal species with the direct-interaction-only model performing as well as the HOI-inclusive model. When HOIs were included during simulation, however, *M. bancana* exhibited a clear rank swap in diameter especially under the low recruitment scenario as it responded to changes in the size of their HOI-sensitive neighbours. When initial recruitment was high, the fast-growing *M. bancana*, as well as *P. polystachya* and *G. nervosa*, exhibited less rank reduction in diameter, which could be important for these species to survive under densely-recruited canopy gaps in the long run. That these species-level rank swaps happened frequently—even when community-level stem size distributions remain qualitatively unaffected—further highlights that ignoring HOIs could lead to very different predictions of community structure. Our results suggest that predictions with and without HOIs are more likely to diverge when individuals are growing more rapidly, as in our low recruitment scenario with lower competition initially, or for some highly-productive systems with sudden resource fluxes. As basal areas build up rapidly at the beginning of gap succession, light depletion often happens more rapidly and the nonlinear size-growth responses of tree species constantly regulate interaction strengths among themselves.

The context-dependence of HOIs begs the question ‘what is the right scale to test for HOIs?’ Seemingly weak HOI-effects on the *instantaneous* diameter growth rates of HOI-insensitive species can accumulate and become long-lasting when *integrated* over the longer lifecycle of perennial plant species, as well as over larger spatial extents where indirect effects of intermediary species domino through direct neighbours via interaction chains. It follows that a *neighbours’ response to* biotic interactions can be just as important as the focal’s response, because the former continuously determines the neighbours’ size and hence their cumulative *effects on* the focal species. Indirect interactions therefore challenge how we conceptualise a focal individual’s biotic milieu or interaction radius: does it extend from a single spatial point as in many pairwise-interaction studies (e.g. Uriarte et al. 2004; Adler, Ellner, and Levine 2010; Comita et al. 2010) or should it be a larger area that includes the neighbours’ neighbours and their decaying yet percolating effects on the focal individual? Due to a lack of spatial data, we were unable to address these questions explicitly here but highlight them as important for future studies. We also acknowledge that without delimiting or estimating the interaction radii (as in Uriarte et al. 2004; Comita et al. 2010; Li et al. 2020) some errors could have entered our parameter estimation (Detto et al. 2019). Future studies should be aware that HOIs—as well as direct interactions—may be common but are simply too weak to be detected (Abrams 1983; Billick and Case 1994; Kleinhesselink, Kraft, and Levine 2019), especially over very short time-scales, when neighbour densities are measured inaccurately (Detto et al. 2019), or if HOIs are only important at a certain life stage.

Due to limited data, we only examined one of the many vital rates impacting perennial plants: size growth. We also lack the data to examine how the effects of biotic interactions on such a single vital rate carry over to influence the final reproductive fitness and hence per capita population growth—the key variable of modern coexistence theory. Other vital rates (e.g. survival and reproduction) that contribute unequally to per capita population growth (Moll and Brown 2008; Adler et al. 2014; Visser et al. 2016) can offset the strong biotic effects on size growth (Broekman et al. 2019). A stronger test of coexistence demands the quantification of the relative contribution of direct and higher-order interactions to multiple vital rates across life stages, and then the estimation of net effects of these biotic interactions on per capita population growth using tools such as population integral projection models (e.g. Chu and Adler 2015). Knowing the effects of neighbours on other vital rates will also improve our diameter-growth simulation (or any other simulation of community dynamics) by incorporating mortality and recruitment.

As the size growth of perennial plants is not only density dependent but also size dependent, another approach is to allow biotic interactions to not only influence the initial diameter growth rate (i.e. the parameter *a* as in this study) but also the ontogenetic effect of size on growth (i.e. the parameter *c* in Equation 1). The latter allows one to test if larger-sized individuals are less sensitive to biotic interactions, and if such a size-conferred storage effect is important for stabilising size-structured communities (Warner and Chesson 1985; Kohyama 1993). Doing so, however, not only further increases the number of parameters in a model that is already data-hungry, but also demands more data collection from larger individuals that are inherently rare in the field (Needham et al. 2018). Although it is possible to circumvent this problem by calculating neighbour basal area from taller neighbour individuals only (e.g. Coomes and Allen 2007), we did not do so because (i) the different height–diameter allometry among species means that a larger-diameter neighbour is not always taller and (ii) defining a taller indirect neighbour is not straightforward (i.e. taller than the focal, the direct neighbour, or both?) Facing the dilemma between trying to fully capture the interplay between size-dependence and density-dependence while keeping the question statistically and logistically tractable, a solution may be to select only a few interaction coefficients that are non-zero with different statistical approaches and biological foresight (Martyn et al. 2021). In this and many previous studies (including those that lumped species into conspecifics versus heterospecifics), the decision to include or not HOIs has been treated as an ‘all or none’ question, tantamount to assuming that HOIs from all neighbour species are either equally important or equally unimportant (Martyn et al. 2021). Although there have been attempts to identify important neighbour species by fitting numerous nested models varying in the identity and number of neighbour species (e.g. Mayfield and Stouffer 2017), implementing this in the Bayesian framework can be computationally impractical. The advancing field of Bayesian variable or model selection (Tenan et al. 2014) can be a good place to start looking for a solution to relax this biologically irrational ‘all or none’ assumption.

## Conclusion

We showed that HOIs are a non-negligible phenomenon at the community level in a tropical forest and an important predictor of diameter growth for a subset of focal tree species. Our study represents one of the early attempts to test for HOIs in perennial plant systems (see also Li et al. 2020). With a high number of HOI parameters that increase exponentially with species number, we expected the effects of most HOIs to be small if not undetectable. Yet we detected the presence of HOIs even with a relatively small dataset, suggesting that many larger datasets can reveal more conclusively the prevalence, direction, and magnitude of HOIs in perennial systems. Though our small dataset limited us to a handful of focal species, the fact that these focal species are all common implies that HOIs are necessarily a widespread phenomenon experienced by many tree *individuals* across the landscape, even if HOIs turn out to be ‘unimportant’ to a large number of rarer species. Last but not least, the empirical quantification of HOIs can only inform us so much about the where and when of biological non-additivity. Much is left to be discovered about the mechanistic why and how of this emergent phenomenon in multi-species communities.

## Author contributions

HRL and DBS conceived of the research idea; HRL, KYC, and ATKY collected the data; HRL analysed the data and led the writing; all authors discussed the results and commented on the manuscript.

## Acknowledgements

We thank Louise Neo, Jolyn W. Loh, Wei Wei Seah, Reuben C.J. Lim, Jon S.Y. Tan, Weng Ngai Lam, Choon Yen Koh and Hugh T.W. Tan for intellectual and field assistance; the National Parks Board of Singapore for granting access (permit NP/RP11-021) and Wildlife Reserves Singapore for funding (Third Ah Meng Memorial Conservation Fund R-154-000-615-720); members of the Stouffer Lab for proofreading. This work was supported by the Marsden Fund Council from New Zealand Government funding, which is managed by the Royal Society Te Apārangi (Grant 16-UOC-008 awarded to DBS). The authors declare no conflict of interest.

## Data availability

Our data will be archived on GitHub upon acceptance.

## Supporting information

**Table S1:** Focal species information.

**Table S2:** Focal species initial recruitment at the beginning of diameter-growth simulation.

**Figure S1:** Interaction coefficients including non-focal neighbour species.

**Figure S2:** Distribution of individual neighbours’ and their cumulative effects.

**Figure S3:** Diameter simulation result for individual trees.

