## Supporting Information for "Non-additive biotic interactions improve predictions of tropical tree growth and impact community size structure"

**Table S1:** The number of individuals and abbreviation of each focal species used in this study.

| Abbreviation | Species | *N* |
|---|---|---|
| ARCHCL | *Archidendron clypearia* | 412 |
| CALOWA | *Calophyllum wallichianum var.* incrassatum | 240 |
| ELAEMA | *Elaeocarpus mastersii* | 131 |
| GARCPA | *Garcinia parvifolia* | 956 |
| GIRONE | *Gironniera nervosa* | 333 |
| MACABA | *Macaranga bancana* | 123 |
| PALAOB | *Palaquium obovatum* | 201 |
| PRUNPO | *Prunus polystachya* | 416 |
| SYZYBO | *Syzygium borneense* | 116 |
| TIMOWA | *Timonius wallichianus* | 340 |

**Table S2:** Initial abundance of focal species under low, median, and high recruitment scenarios at the beginning of the simulation of diameter growth.

| Species | Low | Median | High |
|---|---|---|---|
| *Archidendron clypearia* | 3 | 5 | 10 |
| *Calophyllum wallichianum* var. *incrassatum* | 1 | 1 | 2 |
| *Elaeocarpus mastersii* | 1 | 1 | 2 |
| *Garcinia parvifolia* | 3 | 6 | 10 |
| *Gironniera nervosa* | 1 | 2 | 3 |
| *Macaranga bancana* | 1 | 2 | 3 |
| *Palaquium obovatum* | 1 | 1 | 2 |
| *Prunus polystachya* | 2 | 4 | 7 |
| *Syzygium borneense* | 1 | 1 | 2 |
| *Timonius wallichianus* | 2 | 3 | 5 |
| Total | 16 | 26 | 46 |

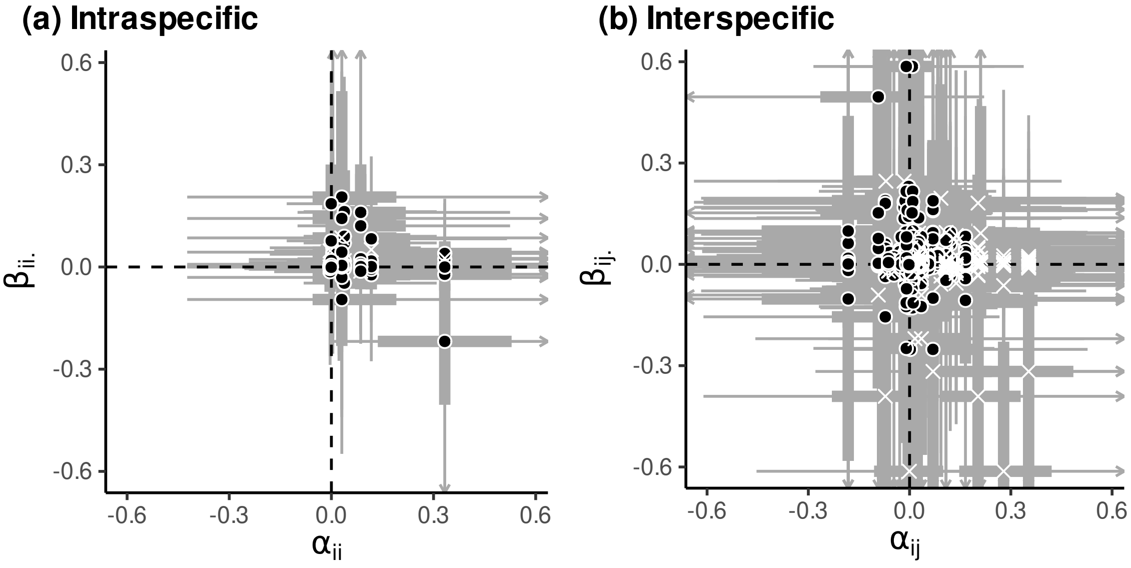


**Figure S1:** The relationship between direct-interaction coefficients ($\alpha$'s) and HOI coefficients ($\beta$'s). This is the same figure as Fig. 1 in the main text, except here it shows the interaction coefficients of non-focal direct and indirect neighbour species as well (white crosses), i.e. $j$ or $k = 11$. Both axes are standardised coefficients that have comparable magnitudes. In **(a)**, intraspecific direct-interaction coefficients ($\alpha_{ii}$) are plotted with their corresponding HOI coefficients ($\beta_{iii}$ or $\frac{1}{2}\beta_{iij}$, together denoted $\beta_{ii\cdot}$). Similarly in **(b)**, interspecific direct-interaction coefficients ($\alpha_{ij}$) are plotted with their corresponding HOI coefficients ($\frac{1}{2}\beta_{iji}$, $\beta_{ijj}$ or $\beta_{ijk}$, together denoted $\beta_{ij\cdot}$). Points and crosses are median estimates with 50% and 95%-tile intervals across the posteriors (thick and thin bars; arrows denote 95%-tile intervals that extend beyond the plot limits).

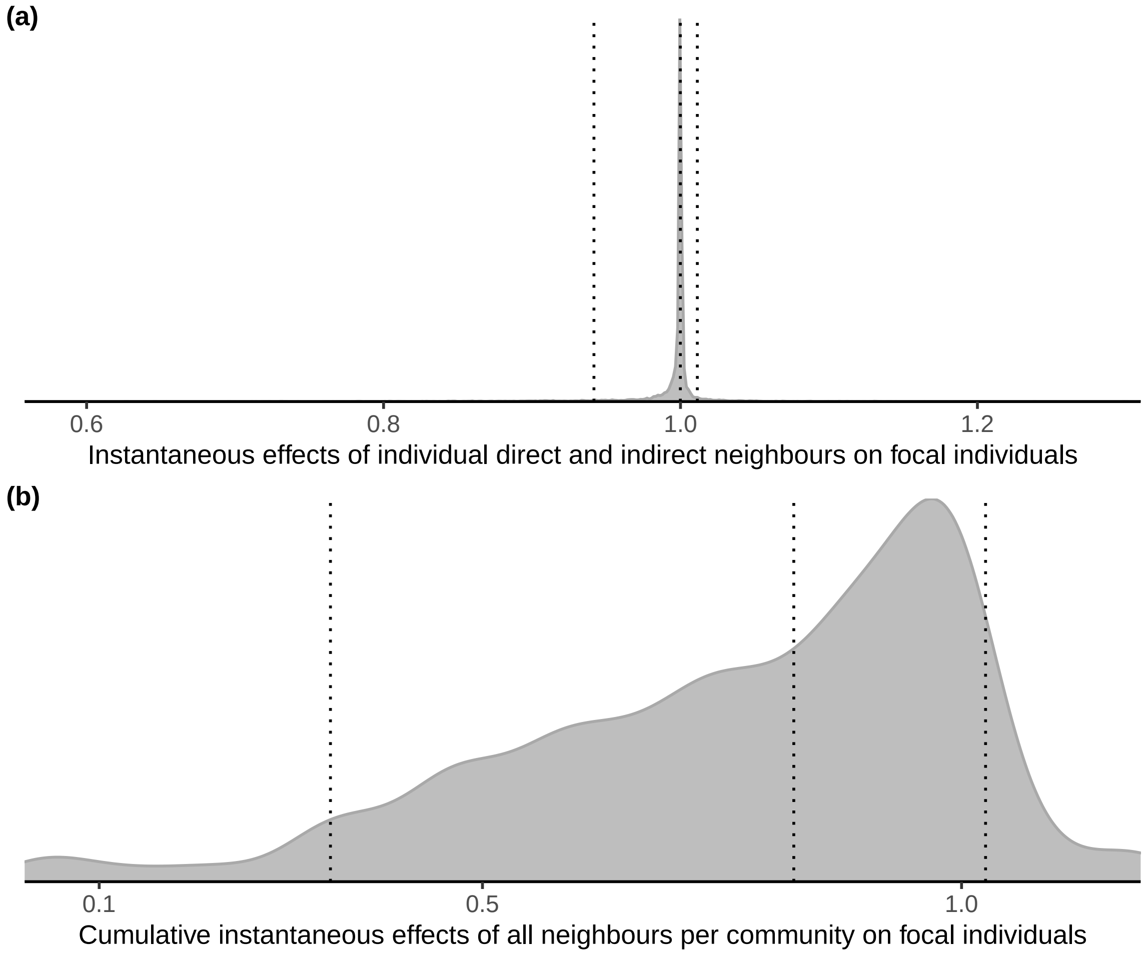


**Figure S2:** The density distribution of **(a)** the instantaneous effects of individual neighbours on focal individuals, i.e., each $\exp(-\alpha_i.A._{,p,q})$ or $\exp(-\beta_i..A._{,p,q}A._{,p,q})$, and **(b)** the cumulative instantaneous effects of all neighbours on focal individuals in a community, i.e., $\exp(-\sum \alpha_i.A._{,p,q})$ or $\exp(-\sum \beta_i..A._{,p,q}A._{,p,q})$. These values represent the proportional change in the average absolute diameter growth rate of a focal individual: the growth rate of a focal individual decreases when effects are $< 1$ ('competitive') and increases when effects are $> 1$ ('facilitative'). Vertical dotted lines denote the 5th percentile, median, and 95th percentile respectively, such that 90% of the data lie between the lower and upper percentiles. The plot limits are bound by the 99%-tile intervals for visual clarity. Note the difference scales on the X-axes.

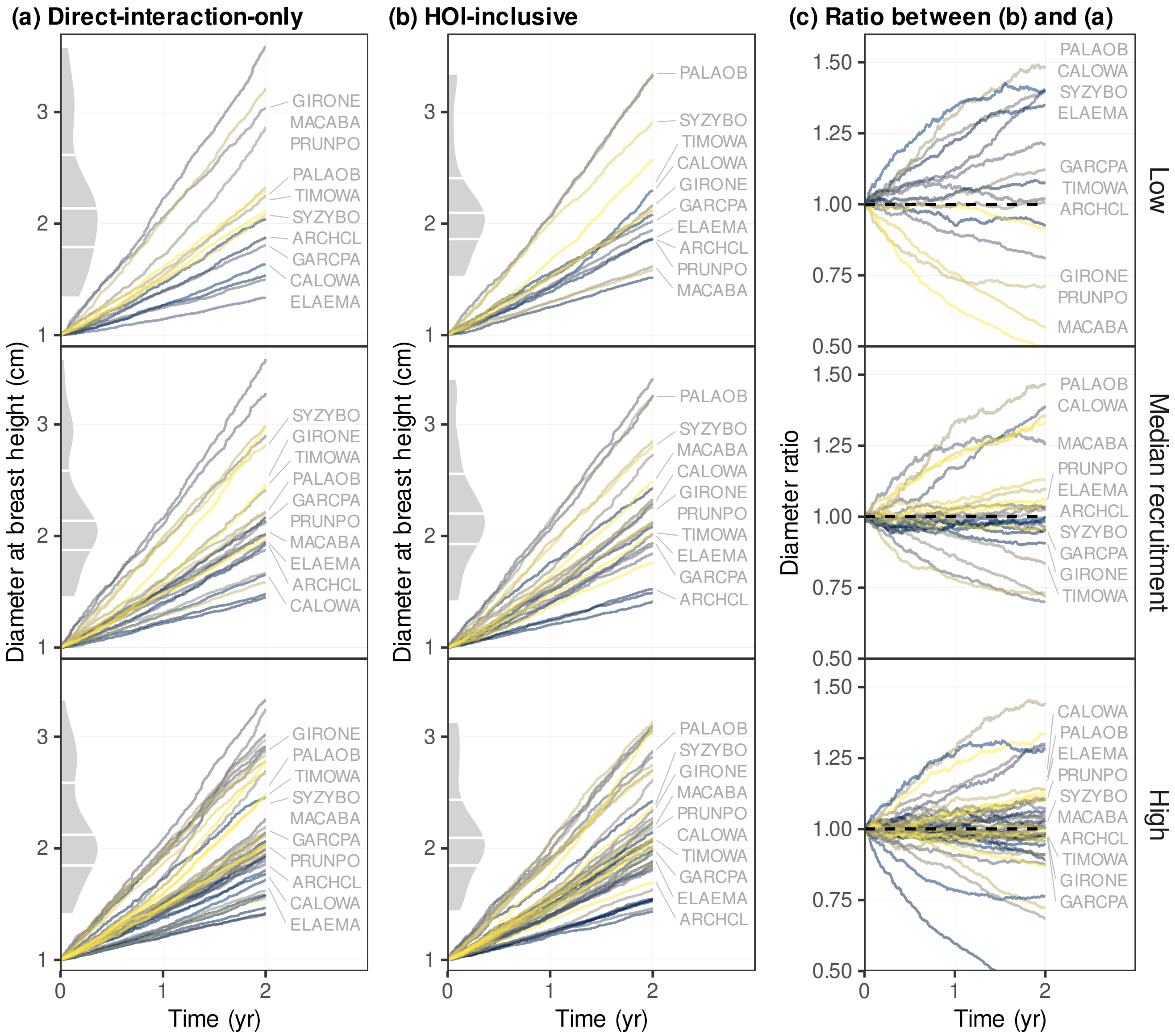


**Figure S3:** The same figure as Fig. 5 in the main text, except that the lines in all panels here are the two-year diameter-growth simulations of *individual trees*, rather than species medians, growing under low, median, and high recruitment scenarios: **(a)** predictions from the direct-interaction-only, **(b)** predictions from the HOI-inclusive model, and **(c)** the species-median ratio of simulated diameter between HOI-inclusive and direct-interaction-only models. Each line represent an individual tree colour-coded per species with yellow to blue representing increasing initial growth rates at small diameters, $a_i'$ (Equation 4 in the main text). Species labels on the right mark the median value of each species, as in Fig. 5 in the main text. Density plots with quartile lines along the Y-axes show the size distribution of individuals trees at year two.
